# Testing the diffusion limitation hypothesis for declining methane uptake in forest soils

**DOI:** 10.64898/2026.03.12.711040

**Authors:** Victor Edmonds

## Abstract

Upland forest soils oxidize 22–38 Tg CH_4_ yr^−1^, roughly 5% of the total atmospheric methane sink. A recent study documented a 53–89% reduction at two long-term ecological research networks in the northeastern United States and attributed it to increased precipitation via diffusion limitation. We tested five predictions of that hypothesis against 27 years of chamber flux data from the Baltimore Ecosystem Study (BES, 1998–2025; *n* = 9,359) and 14 years from the Hubbard Brook Experimental Forest (HBR, 2002–2015).

Four predictions were not supported. At the individual-measurement scale, neither monthly precipitation nor direct soil moisture explained more than 1% of CH_4_ flux variance (*R*^2^ = 0.0008 and 0.0055). While precipitation emerged as a significant interannual predictor when data were aggregated to annual-site means (*β* = 0.249, *p* = 0.002), it did not eliminate the residual multi-decadal decline (*β*_year_ = 0.211, *p* = 0.007). No seasonal moisture–flux structure matched diffusion predictions. Urban and rural BES forests diverged despite sharing a regional precipitation regime (Year×LandUse interaction, *p* = 0.007), and a residual temporal trend persisted after controlling for moisture, temperature, and spatial pseudoreplication (*p* = 0.002). A structural breakpoint at 2002 (BES) and a putative shift at 2011 (HBR) aligned with atmospheric deposition trends rather than precipitation. A fifth test, the Hubbard Brook calcium amendment, yielded a null result that does not discriminate between mechanisms but constrains methanotrophic recovery potential.

These results suggest that precipitation-driven diffusion limitation does not adequately account for the multi-decadal loss of CH_4_ uptake at these sites and point toward chronic biological degradation, potentially through nitrogen-mediated inhibition of high-affinity methanotrophy compounded by structural changes from invasive earthworm activity.

## 1 Introduction

Upland forest soils are a globally significant sink for atmospheric methane. Methanotrophic bacteria in the upper soil horizons oxidize CH_4_ at ambient concentrations (∼1.9 ppm), consuming an estimated 22–38 Tg CH_4_ yr^−1^ (Dutaur and Verchot, 2007; Saunois et al., 2020). This biological sink offsets roughly 5% of total atmospheric CH_4_ removal, but it has been declining.

Ni and Groffman (2018b) documented a 53–89% reduction in CH_4_ uptake across two independent long-term ecological research networks: the Baltimore Ecosystem Study (BES) in Maryland and the Hubbard Brook Experimental Forest (HBR) in New Hampshire. Their analysis of 317 peer-reviewed articles suggested a global decline of 77% over three decades. They attributed the decline to increased precipitation: wetter soils reduce gas-phase diffusivity, physically limiting the supply of atmospheric CH_4_ to methanotrophs in the soil column.

This attribution rests on the diffusion limitation hypothesis, which holds that soil moisture is the primary rate-limiting control on CH_4_ uptake (Striegl, 1993; Ball et al., 1997; Ridgwell et al., 1999). The mechanism is intuitive: as rainfall fills soil pore spaces, the volume available for gas-phase transport shrinks, and because atmospheric CH_4_ must diffuse from the surface to the methanotrophs below, reduced air-filled porosity directly limits substrate supply. The framework is well established. Multiple global models parameterize the soil CH_4_ sink through diffusion-based equations (Del Grosso et al., 2000), and experimental manipulations at Harvard Forest have demonstrated that throughfall exclusion increases CH_4_ uptake while snow cover suppresses it (Borken et al., 2006). The hypothesis is mechanistically coherent and has been the default explanation for temporal variation in forest CH_4_ fluxes for three decades.

The organisms responsible for this oxidation are primarily uncultured high-affinity methanotrophs, principally the Upland Soil Cluster alpha (USC*α*) (Kolb, 2009; Knief, 2015). Unlike the low-affinity methanotrophs active at the elevated concentrations found in wetlands and landfills, USC*α* specialize in atmospheric-concentration oxidation. They are extremely slow-growing, with recovery timescales measured in decades to centuries after disturbance (Lim et al., 2024). This biology matters for evaluating the diffusion hypothesis: any mechanism that degrades the methanotrophic community, rather than merely restricting gas supply, would produce a decline that persists independently of moisture conditions.

The diffusion hypothesis also makes specific predictions. If diffusion limitation controls the long-term decline in CH_4_ uptake, then: (1) soil moisture should explain a substantial fraction of flux variance; (2) the moisture–flux relationship should exhibit seasonal structure consistent with site-specific precipitation regimes; (3) sites receiving identical precipitation should exhibit identical flux trajectories regardless of soil chemistry; (4) flux recovery should be general across sites sharing a precipitation regime; and (5) after controlling for moisture, no residual temporal trend should remain. A parallel, non-climatic hypothesis (physical destruction of methanotrophic habitat by invasive earthworms) makes overlapping predictions with the biological alternatives considered here and is evaluated in the Discussion.

Ni and Groffman (2018b) tested the association between precipitation trends and CH_4_ decline using data through 2016. They did not test direct soil moisture measurements against flux, apply formal changepoint detection, or evaluate the calcium amendment experiment at Hubbard Brook as a natural manipulation. Direct in-situ volumetric water content (VWC) data from BES soil sensors (2011–2020) were not incorporated.

We extend the BES record by 9 years (through 2025), add direct VWC measurements, and apply five independent tests to the diffusion hypothesis. The analysis incorporates over 9,000 quality-controlled flux measurements, PRISM climate data, NADP wet deposition records, lysimeter soil solution chemistry, and vegetation surveys. Four of the five predictions were not supported; the fifth (a calcium amendment experiment) yields a null result consistent with diffusion limitation but also with biological alternatives. Taken together, the results suggest that precipitation-driven diffusion limitation, as applied to these two LTER networks, does not adequately account for the observed decline.

## 2 Methods

### 2.1 Study sites

The Baltimore Ecosystem Study (BES) is an urban-to-rural Long Term Ecological Research site in the greater Baltimore, Maryland metropolitan area (39.3°N, 76.6°W). CH_4_ flux measurements span nine forest sites along an urbanization gradient, from urban parks (Gwynns Falls, Hillsdale) to exurban forests (Oregon Ridge). The Hubbard Brook Experimental Forest (HBR) is located in the White Mountains of New Hampshire (43.9°N, 71.7°W). CH_4_ flux data are available from Watershed 1 (WS1), which received a calcium silicate amendment in October 1999, and the reference Watershed 6 at Bear Brook (WS6-BB). Site characteristics are summarized in Table 1.

**Table 1.** Study site characteristics.

| Site | Network | Setting | Latitude | Forest type | Elev. (m) | Period |
| --- | --- | --- | --- | --- | --- | --- |
| Hillsdale (HD) | BES | Urban | 39.35°N | Mixed deciduous | 60 | 1998–2025 |
| Leakin Park (LEA) | BES | Urban | 39.32°N | Mixed deciduous | 70 | 1998–2025 |
| Oregon Ridge (ORM) | BES | Rural | 39.50°N | Mixed deciduous | 210 | 1998–2025 |
| Oregon Ridge Upper (ORU) | BES | Rural | 39.50°N | Mixed deciduous | 240 | 1998–2025 |
| WS1 (Ca-treated) | HBR | Montane | 43.94°N | Northern hardwood | 500 | 2002–2015 |
| WS6-BB (Reference) | HBR | Montane | 43.94°N | Northern hardwood | 530 | 2002–2015 |

*Note:* BES includes additional sites (MCD, GLY, UMBC, CAH) used in pooled analyses. GB and ORLR were excluded (see Sect. 2.3). The pooled BES analytical dataset comprises 9,359 measurements after quality control (Sect. 2.3); site-level counts are reported in the analysis code.

### 2.2 Data sources

#### CH_4_ flux

BES trace gas collection data (1998–2025) were obtained from the BES LTER data archive (knb-lter-bes). Measurements were made using static chambers with gas chromatography. The dataset contains 10,674 individual flux measurements across all sites and years. HBR CH_4_ flux data (2002–2015) were obtained from the Hubbard Brook data archive (knb-lter-hbr.207), comprising 231 annual and 1,764 monthly observations.

#### Climate

Monthly precipitation and mean temperature for each site were obtained from the PRISM Climate Group (Oregon State University) at 4 km resolution. Separate extractions were performed for BES and HBR site coordinates.

#### Soil moisture

Hourly volumetric water content (VWC) measurements from BES soil sensor networks (2011–2020) were obtained from the BES data archive (knb-lter-bes.3400). The dataset contains 691,965 hourly readings from nine sensor locations across five sites.

#### Atmospheric deposition

Wet deposition of sulfate 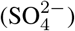 and inorganic nitrogen 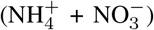 were obtained from the National Atmospheric Deposition Program (NADP) National Trends Network. Station NH02 (Hubbard Brook) provided sulfate records from 1978–2023. Station MD99 (Beltsville, MD) provided inorganic nitrogen records from 1999–2023.

#### Soil solution chemistry

Lysimeter 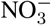 concentrations (1999–2025) were obtained from the BES data archive (knb-lter-bes.428), comprising 20,479 measurements across three forested sites (Hillsdale, Leakin Park, Oregon Ridge).

#### Soil properties

Physical, chemical, and biological properties of forest soils at 0–10 cm depth were obtained from knblter-bes.584 for four BES sites (Hillsdale, Leakin Park, Oregon Ridge, Oregon Ridge Upper). Properties include C:N ratio, microbial biomass carbon, net nitrogen mineralization, and net nitrification rates.

#### Vegetation

Seedling cover data across three BES census years (1998, 2003, 2015) were obtained from knb-lter-bes.3300 for three forested sites.

### 2.3 Data quality control

Two BES sites were excluded from all sink analyses: Gwynns Falls (GB), an urban stream-adjacent site, and Oregon Ridge Lower Riparian (ORLR). Together these sites produced 63 extreme positive CH_4_ measurements (fluxes up to 7,234 mg C m^−2^ h^−1^) representing ebullition from saturated riparian soils rather than the upland forest sink. These measurements comprised 0.6% of the total dataset.

An additional per-measurement filter removed values exceeding *±*3 standard deviations within each site-year group. This two-stage filter (hotspot site exclusion followed by within-group trimming) reduced the analytical dataset from 10,674 to 9,359 measurements. The filter criteria, site exclusion rationale, and the number of observations removed at each stage are documented in the analysis code.

### 2.4 Prediction 1: Moisture–flux regressions

#### Precipitation proxy (climatological test)

For each BES flux measurement, we matched the corresponding monthly total precipitation from PRISM. Ordinary least-squares regression of CH_4_ flux on monthly precipitation was performed on the pooled dataset (*n* = 9,359). This tests the diffusion hypothesis as Ni and Groffman (2018b) originally framed it: using precipitation as a proxy for soil moisture at the climatological scale. The diffusion hypothesis predicts a significant negative relationship (wetter months, less uptake).

#### Direct soil moisture (mechanistic test)

BES VWC sensor readings were matched to CH_4_ flux measurements by site and date. Where multiple hourly VWC readings existed for a measurement day, the daily mean VWC was used. This yielded 2,415 paired VWC–flux observations (2011–2020). OLS regression was performed as above. Because gas-phase diffusivity (*D*_*p*_) is physically controlled by air-filled porosity—the fraction of soil pore space not occupied by water—precipitation is an indirect proxy at best. A given rainfall event may or may not change soil moisture depending on antecedent conditions, drainage, and evapotranspiration. The VWC test therefore provides the mechanistically appropriate evaluation of diffusion limitation, while the precipitation test evaluates the hypothesis as Ni and Groffman (2018b) originally framed it. A significant VWC–flux relationship would support diffusion limitation even if precipitation proved to be a poor proxy.

### 2.5 Prediction 2: Seasonal stratification

The dataset was stratified by meteorological season (Winter: DJF; Spring: MAM; Summer: JJA; Fall: SON). Separate precipitation– flux regressions were performed within each season. In the humid mid-Atlantic climate of Baltimore, summer receives the highest total precipitation, but summer evapotranspiration (ET) from the deciduous canopy is also at its annual maximum, making summer soils the *driest* of the year in terms of water-filled pore space. Diffusion theory therefore predicts that summer should exhibit the *highest* methane uptake (maximum air-filled porosity) and the weakest precipitation–flux coupling (because precipitation inputs are rapidly offset by ET). The strongest moisture–flux coupling should occur in spring or fall, when precipitation exceeds ET and soils are wettest. We tested whether any seasonal *R*^2^ exceeded a negligible threshold and whether the seasonal flux pattern tracked the expected soil moisture seasonality.

### 2.6 Prediction 3: Calcium amendment experiment

The Hubbard Brook WS1 calcium silicate experiment provides a natural manipulation in which soil chemistry was altered while precipitation remained identical. WS1 (treated, October 1999) and WS6-BB (reference) occupy adjacent watersheds at similar elevation and aspect. Both receive the same precipitation inputs.

We compared mean CH_4_ flux between WS1 and WS6-BB and computed Cohen’s *d* as the standardized effect size. We also computed the frequency of source events (positive CH_4_ flux, indicating net emission) at each site. The diffusion hypothesis predicts identical flux at both sites (same rain, same response), so a null result is expected under diffusion limitation. However, the comparison also tests whether reversing soil acidification—a major biogeochemical consequence of the calcium addition— restores CH_4_ uptake capacity. A null result therefore constrains the biological interpretation by establishing that 14 years of improved soil chemistry did not detectably alter the methanotrophic sink.

### 2.7 Prediction 4: Urban–rural divergence

BES sites were classified as urban (Hillsdale, Leakin Park) or rural (Oregon Ridge, Oregon Ridge Upper) following the BES urbanization gradient. Post-2012 linear trends in annual CH_4_ flux were computed separately for each group. The cutoff year (2012) was chosen based on Ni and Groffman (2018b)’s observation of possible recovery. Under a simple diffusion framework, sites sharing a regional precipitation regime should exhibit broadly convergent flux trajectories. However, urban and rural soils may differ in bulk density, macropore structure, and compaction history, so identical precipitation does not necessarily produce identical air-filled porosity. Divergent trajectories therefore provide suggestive but not definitive evidence for a non-climatic driver.

### 2.8 Prediction 5: Multi-predictor model

Two models were fit to the measurement-scale data. The primary model was a linear mixed-effects model (LMM) with a random intercept for site, CH_4_ ∼ Precipitation + Temperature + Year + (1 | Site), fit via restricted maximum likelihood (REML). The random intercept absorbs systematic differences between sites (e.g., one site consistently producing higher uptake than another), ensuring that fixed-effect estimates reflect within-site temporal relationships rather than between-site contrasts. This accounts for pseudoreplication from repeated chamber measurements at fixed locations. For transparency, an ordinary leastsquares (OLS) model was also fit for comparison. All predictors were standardized (*z*-scored) to allow direct comparison of coefficient magnitudes. The diffusion hypothesis predicts that after controlling for moisture, no significant temporal trend (Year effect) should remain.

To address the possibility that chamber-level measurement noise obscures a real moisture signal, the OLS model was additionally fit at three temporal scales: (1) individual measurements (*n* = 9,359), (2) seasonal-site means aggregated by site × season × year (*n* = 625), and (3) annual-site means aggregated by site × year (*n* = 169). If diffusion limitation operates at longer timescales than individual measurements can resolve, aggregation should increase the precipitation signal relative to Year. The diffusion hypothesis predicts that precipitation should dominate and Year should become non-significant at the aggregated scales.

### 2.9 Changepoint detection

To identify years in which the CH_4_ flux time series underwent abrupt structural shifts, as opposed to gradual trends, we applied the Pruned Exact Linear Time (PELT) changepoint algorithm (Killick et al., 2012). PELT partitions a time series into segments that minimize within-segment variance, identifying the points where the statistical properties of the data shift. It was applied to annual median CH_4_ flux at BES and HBR independently. Annual medians were used rather than means because extreme positive events (even after outlier filtering) distort the mean, while the median tracks the bulk behavior of the soil sink. Penalty calibration was performed through sensitivity analysis across a range of penalty values (0.01–10). A penalty of 0.1 produced stable, interpretable breakpoints at both sites. The sensitivity analysis is documented in the analysis code.

### 2.10 Deposition and soil chemistry context

NADP wet deposition data were plotted against CH_4_ flux time series to assess temporal alignment between deposition trends and flux breakpoints. Lysimeter NO^−^_3_ concentrations were analyzed by site for monotonic trends using OLS regression against time.

### 2.11 Statistical considerations

The 9,359 individual flux measurements are not fully independent. Repeated measurements from fixed chamber locations introduce spatial pseudoreplication, and temporal autocorrelation is expected in long environmental time series. We address these concerns in four ways. First, all primary results are reported at both the individual-measurement and site-year aggregated scales. Second, changepoint detection (PELT) is applied to annual time series rather than raw measurements, eliminating within-year autocorrelation. Third, the five tests are independent of one another; the conclusion rests on the convergence of multiple lines of evidence, not on the statistical significance of any single test. Fourth, the multi-predictor model (Prediction 5) is fit as a linear mixed-effects model (LMM) at two levels of spatial nesting: a random intercept for site and a random intercept for individual collar (Site×Plot×Chamber); the collar-level LMM serves as the primary analysis, with the site-level LMM and OLS providing comparisons. Residuals are checked for temporal autocorrelation via Durbin-Watson statistics.

All analyses were performed in Python 3.x using pandas, numpy, scipy, statsmodels, matplotlib, seaborn, and ruptures. The complete analysis is implemented in a single reproducible script (`master_analysis.py`) with portable relative paths. All data are publicly available from the respective LTER data archives.

## 3 Results

### 3.1 Prediction 1: Moisture–flux regressions

Monthly precipitation explained 0.08% of the variance in CH_4_ flux across the pooled BES dataset (*R*^2^ = 0.0008, slope = 0.0011, *p* = 0.007, *n* = 9,359; Fig. 1a). The slope was positive (more precipitation associated with less negative flux, i.e., reduced uptake), consistent with the predicted direction under diffusion limitation, but the magnitude is negligible. A temporal scale mismatch is inherent to this test: each chamber measurement captures a single hour of gas exchange, while monthly precipitation integrates 30 days of rainfall. This mismatch weakens the expected correlation even if diffusion limitation operates on shorter timescales, making the precipitation test a conservative evaluation of the hypothesis.

**Figure 1.**
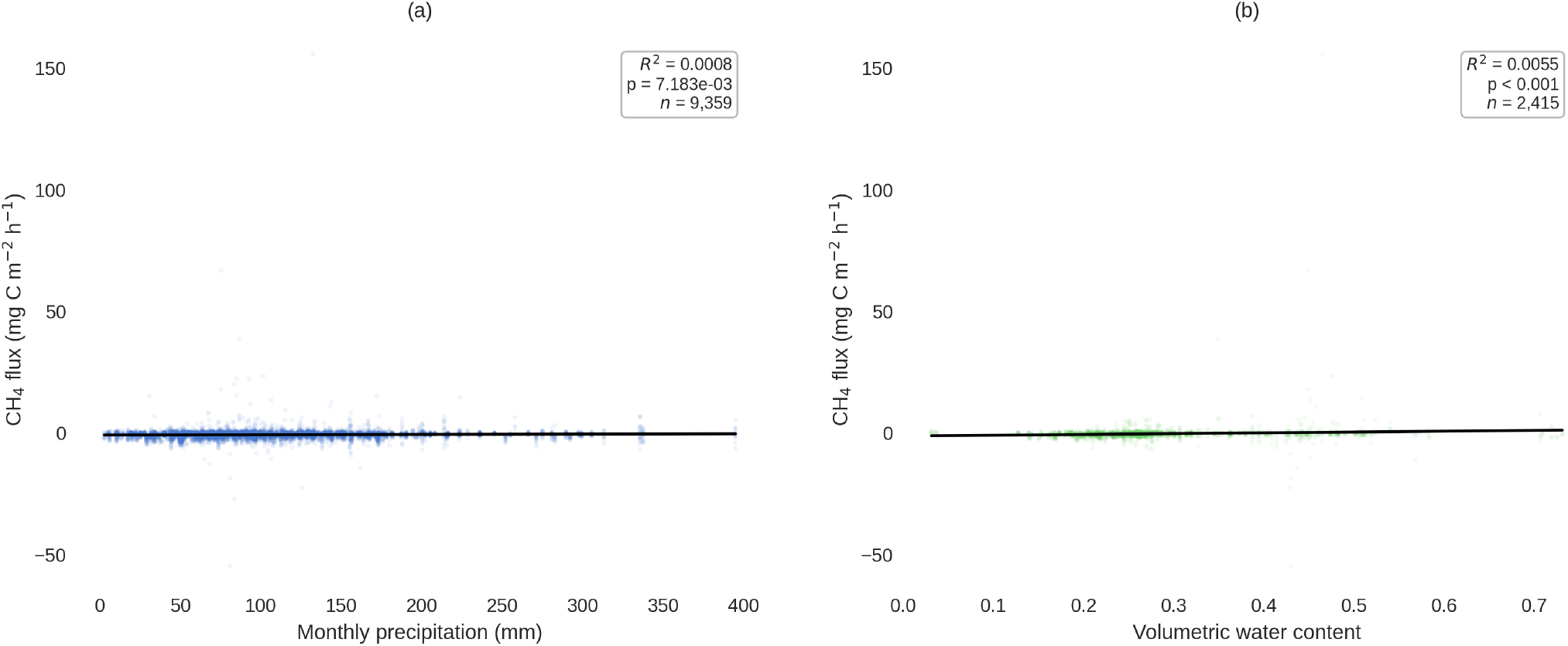
Moisture–flux regressions at BES. (a) Monthly precipitation versus CH_4_ flux, 1998–2025 (*n* = 9,359; *R*^2^ = 0.0008). (b) In situ volumetric water content versus CH_4_ flux, 2011–2020 (*n* = 2,415; *R*^2^ = 0.0055). Neither the climatological proxy nor the direct mechanistic variable explains more than 1 % of flux variance.

Direct in-situ VWC performed marginally better but remained negligible: *R*^2^ = 0.0055 (slope = 3.26, *p <* 0.001, *n* = 2,415; Fig. 1b). The direct soil moisture measurement, which the diffusion hypothesis identifies as the mechanistically relevant variable, explained less than 1% of flux variance over the 2011–2020 measurement period.

Neither the proxy nor the direct measurement approaches the explanatory power expected of a rate-limiting control (Table 2). Borken et al. (2006) found moisture explained substantial variance at Harvard Forest under experimental manipulation; in observational data spanning 27 years at BES, the relationship is effectively absent.

**Table 2.** Moisture–flux regression results.

| Predictor | Scale | *n* | *R*<sup>2</sup> | Slope | *p* | 95% CI |
| --- | --- | --- | --- | --- | --- | --- |
| Precipitation (mm) | Measurement | 9,359 | 0.0008 | 0.0011 | 0.007 | [0.0003, 0.0020] |
| In-situ VWC | Measurement | 2,415 | 0.0055 | 3.26 | <0.001 | [1.52, 5.00] |
| Precip (seasonal) | Winter | 2,069 | 0.0023 | 0.0014 | 0.031 | [0.0001, 0.0026] |
| Precip (seasonal) | Spring | 2,489 | 0.0072 | 0.0025 | <0.001 | [0.0013, 0.0037] |
| Precip (seasonal) | Summer | 2,501 | 0.0048 | 0.0040 | <0.001 | [0.0018, 0.0063] |
| Precip (seasonal) | Fall | 2,300 | 0.0001 | −0.0002 | 0.70 | [−0.0014, 0.0009] |

A caveat applies to the VWC test: the soil moisture data span 2011–2020, a period that postdates the BES structural breakpoint at 2002 (Sect. 3.6). If the methanotrophic community was already degraded by 2011, the VWC test evaluates moisture– flux coupling in a system where the biological capacity for atmospheric CH_4_ oxidation may have been diminished. The low *R*^2^ is therefore ambiguous: it could indicate that diffusion was never the rate-limiting step, or that diffusion control was present historically but is no longer detectable because the biological sink has weakened. The precipitation–flux test (1998–2025) spans the full record, including the pre-breakpoint period, and also shows negligible explanatory power (*R*^2^ = 0.0008).

As a sensitivity check, re-running the precipitation–flux regression without the *±*3 SD per-site-year filter (retaining 126 extreme measurements) actually *weakened* the precipitation signal: *R*^2^ dropped from 0.0008 to 0.0003 and became nonsignificant (*p* = 0.085). The extreme positive measurements, microsites where saturation-driven methanogenesis overwhelms uptake, do not reinforce a precipitation–flux relationship; they add noise that further obscures any diffusion signal (see Sect. 4.11 for details).

The moisture–flux relationship was not always this weak. Stratifying the BES record at the 2002 PELT changepoint (Sect. 3.6) reveals that pre-breakpoint (1998–2002) precipitation explained 0.57% of flux variance (*n* = 1,250, *R*^2^ = 0.0057, *p* = 0.008); roughly 7× the full-record value. A quadratic model accounting for the unimodal diffusion relationship reached 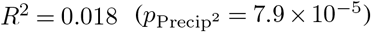. After 2002, the relationship vanished entirely (*R*^2^ = 0.0001, *p* = 0.31). A caveat: the 2002 split point was identified by PELT on annual median fluxes from the same dataset, so this stratification is post-hoc. A permutation test shuffling year labels (*n* = 1,000) yielded *p* = 0.057 for the observed Δ*R*^2^, indicating that the pre/post coupling difference is suggestive but not formally significant at *α* = 0.05 (Sect. 4.11). PELT optimized for a shift in flux *level*, not for moisture–flux *coupling strength*, so the pre/post comparison tests a quantity that was not used to select the breakpoint, but this distinction does not fully resolve the circularity concern. This pattern is consistent with a weak moisture–flux coupling that existed while the methanotrophic community was intact, and that disappeared after the structural break as the community’s capacity to respond to moisture fluctuations degraded. Monthly precipitation is an inherently blunt proxy for the instantaneous soil moisture experienced by a chamber at the time of measurement; the direct VWC test above provides the mechanistically appropriate evaluation, but even this more relevant variable explains less than 1% of variance.

### 3.2 Prediction 2: Seasonal stratification

Seasonal precipitation–flux *R*^2^ values were: Spring 0.0072, Summer 0.0048, Winter 0.0023, Fall 0.0001. No season exceeded 1%. Although summer has the highest total precipitation, high evapotranspiration from the deciduous canopy makes summer soils the driest of the year. Diffusion theory therefore predicts the strongest moisture–flux coupling in spring or fall, when soils are wettest. Spring showed the highest *R*^2^ (0.0072), but this remains negligible. Fall, when soils begin to rewet after the growing season, showed the weakest coupling (*R*^2^ = 0.0001), opposite to diffusion predictions. At no season did precipitation explain a meaningful fraction of flux variance.

### 3.3 Prediction 3: Calcium amendment experiment

Mean CH_4_ flux at WS1 (Ca-treated) was −1.0414 mg C m^−2^ h^−1^. Mean flux at WS6-BB (reference) was −1.0346 mg C m^−2^ h^−1^. Cohen’s *d* = −0.012, a negligible standardized effect (|*d*| *<* 0.2 is conventionally classified as small; Table 3). Both sites declined at the same rate. WS1 had a source event frequency of 0.0%; WS6-BB had 1.8%.

**Table 3.**
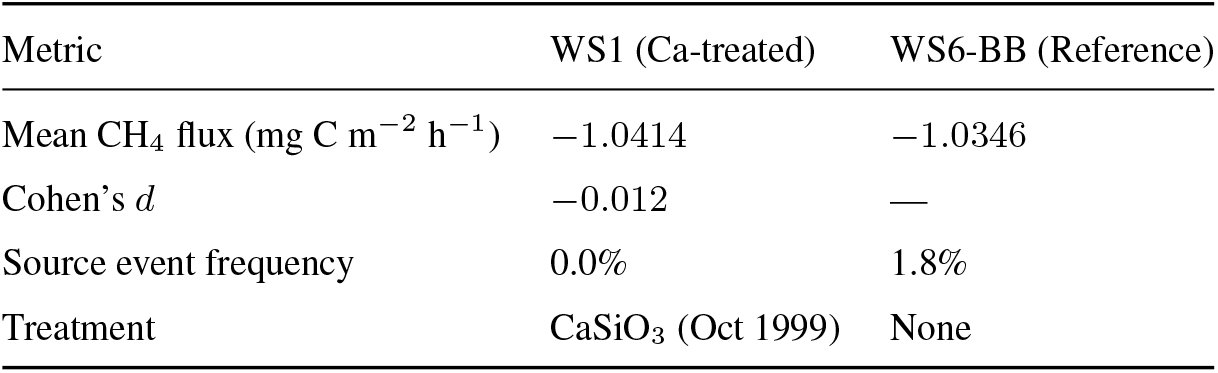
Calcium amendment experiment: WS1 (treated) vs. WS6-BB (reference), Hubbard Brook, 2002–2015.

| Metric | WS1 (Ca-treated) | WS6-BB (Reference) |
| --- | --- | --- |
| Mean CH<sub>4</sub> flux (mg C m<sup>−2</sup> h<sup>−1</sup>) | −1.0414 | −1.0346 |
| Cohen’s *d* | −0.012 | — |
| Source event frequency | 0.0% | 1.8% |
| Treatment | CaSiO<sub>3</sub> (Oct 1999) | None |

At first glance, the null result appears consistent with the diffusion hypothesis, which predicts that a chemical amendment should not affect gas-phase transport. The soils at Hubbard Brook are Spodosols (acidic, leached forest soils with a characteristic iron-enriched B-horizon), derived from glacial till and texturally coarse, sandy loams with very low clay content. Unlike the heavy agricultural soils where liming promotes clay flocculation and macroporosity changes (Haynes and Naidu, 1998), surface wollastonite application to these sandy soils is unlikely to have meaningfully altered bulk density or gas-phase diffusivity in the deeper profile. Identical flux between WS1 and WS6-BB is therefore exactly what the physical diffusion hypothesis predicts, and this test does not discriminate between diffusion and biological mechanisms on physical grounds alone. Its value lies instead in constraining the biological interpretation: the calcium silicate addition reversed soil solution chemistry, increased soil pH, and restored forest growth at WS1 (Battles et al., 2014; Johnson et al., 2014), yet did not detectably alter the CH_4_ sink over 14 years of monitoring. If acidification-mediated damage to the methanotrophic community were readily reversible, recovery should have been apparent at WS1 over this timescale.

If the decline reflected reversible acidification damage to methanotrophs, calcium amendment should have initiated recovery over the 14-year monitoring period, yet no such recovery was observed. This is consistent with either irreversible community degradation (see Lim et al., 2024, who document recovery timescales of decades to centuries for high-affinity methanotrophs) or a mechanism unrelated to soil acidity per se.

However, a stronger interpretation is available. The high-affinity atmospheric methanotrophs (USC*α*) that dominate upland forest soils are adapted to acidic conditions, with some clades explicitly acidophilic (Täumer et al., 2021). Liming raises soil pH and dramatically stimulates nitrification (Groffman et al., 2006), producing 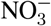 and 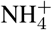 intermediates that competitively inhibit methane monooxygenase (MMO), the enzyme responsible for the initial oxidation step (Schnell and King, 1994). The CaSiO_3_ addition therefore did not merely fail to restore the methanotrophic community—it actively created a biochemically hostile environment for USC*α* by simultaneously raising pH above their acidophilic optimum and flooding the soil with the primary enzymatic inhibitor. The null result is not simply ambiguous between mechanisms; it is the expected outcome of a treatment that inadvertently suppressed the very organisms it would need to recruit. This reinterpretation elevates the calcium amendment from a non-discriminating null result to positive evidence for biological control: the treatment tested a specific biogeochemical pathway and produced the response that pathway predicts.

### 3.4 Prediction 4: Urban–rural flux trajectories

Post-2012 trends in CH_4_ flux diverged between urban and rural BES forests (Fig. 2). Rural forests showed a significant continuing decline (slope = −0.170, *p* = 0.001). Urban forests showed no significant trend (slope = −0.049, *p* = 0.31). However, comparing two separate *p*-values does not formally test whether the trajectories differ (Gelman and Stern, 2006). An interaction model on post-2012 BES chamber data (CH_4_ ∼ Year × LandUse; *n* = 4,639; 2012–2025) confirmed that the divergence is statistically significant: the Year×LandUse interaction was *β* = −0.071 (*p* = 0.007). A secondary analysis using monthly aggregated data (2012–2016; *n* = 234) yielded a consistent result (*β* = −0.122, *p* = 0.015).

**Figure 2.**
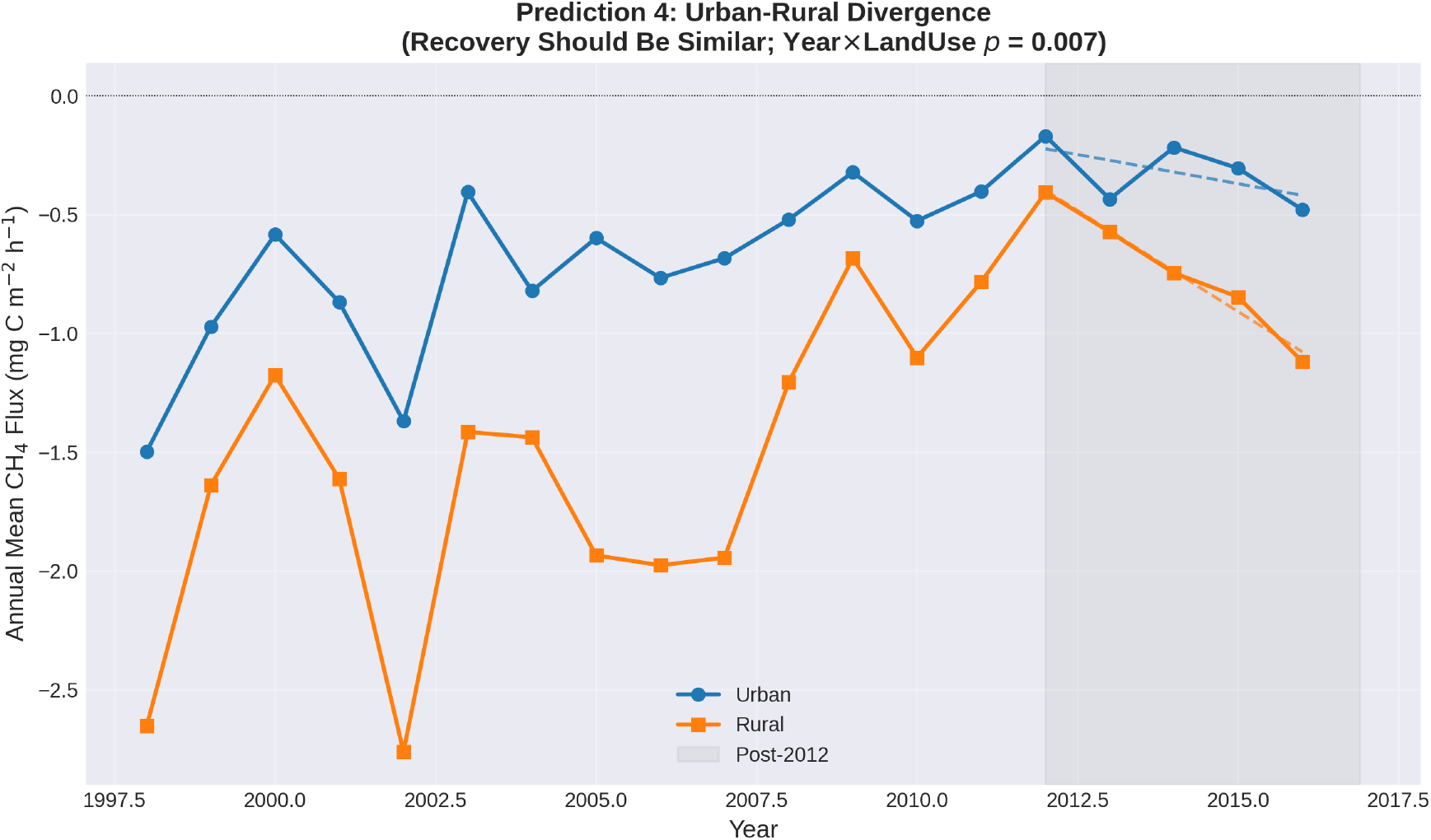
Post-2012 CH_4_ flux trends at urban versus rural BES sites. Rural forests show continuing decline (*p* = 0.001); urban forests show no significant trend (*p* = 0.31). The Year×LandUse interaction is significant (*p* = 0.007), confirming divergent trajectories under a shared precipitation regime.

The two groups share regional precipitation, temperature, and atmospheric CO_2_ concentrations. Under a simple diffusion framework, shared precipitation should produce broadly similar flux trajectories. However, urban and rural soils differ in bulk density, macropore structure, and compaction history, so identical precipitation does not necessarily yield identical air-filled porosity. The divergence is therefore ambiguous with respect to diffusion: it could reflect differences in soil physical properties, or it could implicate non-climatic drivers such as deposition history, fragmentation, heat island effects, or microbial community composition. The pattern is more naturally explained by the latter, but does not by itself rule out a diffusion-mediated contribution. One possibility is that urban forests have reached a functional floor: if the high-affinity methanotrophic community has already been depleted at urban sites, minimal residual oxidation capacity would leave little room for further decline regardless of moisture conditions. This interpretation is evaluated in Sect. 4.8.

### 3.5 Prediction 5: Multi-predictor models across temporal scales

The linear mixed-effects model (LMM) with a random intercept for site yielded standardized coefficients of: Temperature (*β* = −0.184, *p <* 10^−13^), Precipitation (*β* = 0.103, *p <* 10^−4^), and Year (*β* = 0.081, *p* = 0.003). The site random intercept variance (0.40) was modest relative to the residual variance (5.32), confirming that site-level clustering accounts for only a small fraction of total variability. Year remained significant after accounting for the hierarchical data structure, indicating a residual temporal trend that persists after controlling for both moisture and temperature. The diffusion hypothesis predicts that moisture should dominate and Year should become non-significant; this does not occur.

Because chamber measurements are repeated at fixed collar locations within sites, we also fit a collar-level model (random intercept for each of 64 unique Site×Plot×Chamber combinations). The collar-level model was preferred by AIC (ΔAIC = −24 relative to the site-level model). Fixed-effect estimates were essentially unchanged: Year *β* = 0.084 (*p* = 0.002), Precipitation *β* = 0.101 (*p <* 10^−4^), Temperature *β* = −0.181 (*p <* 10^−13^). Accounting for collar-level spatial pseudoreplication slightly *increased* the Year coefficient rather than attenuating it. LMM residuals showed weak temporal autocorrelation (mean per-site lag-1 *r* = 0.19; Durbin-Watson range 1.34–2.04 across sites). This level of autocorrelation may modestly inflate the Year *p*-value but is unlikely to alter its significance at *α* = 0.05. A random-slopes specification (Year|Site) rendered the fixed Year effect non-significant (*β* = 0.016, *p* = 0.81), indicating that the temporal trend varies substantially across sites—consistent with the urban–rural divergence documented in Sect. 3.4. The precipitation coefficient was unchanged (*β* = 0.100), confirming that moisture’s weak explanatory power is insensitive to whether sites are forced to share a common temporal slope (see Sect. 4.11).

For comparison, the OLS model (CH_4_ ∼ Precipitation + Temperature + Year) explained 1.2% of total flux variance (*R*^2^ = 0.012, *F* = 37.82, *p <* 0.001, *n* = 9,359). The OLS Year coefficient (*β* = 0.192) was attenuated in the LMM (*β* = 0.081), indicating that part of the OLS temporal trend reflected site-level differences absorbed by the time term. Precipitation (*β* = 0.104) and Temperature (*β* = −0.167) were essentially unchanged between models (Table 4).

**Table 4.** Multi-predictor model (CH_4_ ∼ Precipitation + Temperature + Year) across temporal scales. All predictors standardized.

| Scale | $n$ | $R^2$ | $\beta_{\text{ppt}}$ | $p_{\text{ppt}}$ | $\beta_{\text{temp}}$ | $p_{\text{temp}}$ | $\beta_{\text{year}}$ | $p_{\text{year}}$ |
| --- | --- | --- | --- | --- | --- | --- | --- | --- |
| Measurement | 9,359 | 0.012 | 0.104 | $4.7 \times 10^{-5}$ | -0.167 | $6.8 \times 10^{-11}$ | 0.192 | $6.4 \times 10^{-15}$ |
| Seasonal-site | 625 | 0.031 | 0.133 | 0.037 | -0.098 | 0.12 | 0.224 | 0.0002 |
| Annual-site | 169 | 0.108 | 0.249 | 0.002 | 0.013 | 0.87 | 0.211 | 0.007 |

In both specifications, precipitation contributed the least explanatory power at the measurement scale.

In the humid mid-Atlantic, temperature covaries positively with precipitation (warm months are also wet). The negative coefficient is consistent with warm-wet conditions suppressing uptake through increased water-filled pore space; it does not independently support diffusion limitation because the temperature–moisture covariance cannot be separated at the measurement scale without continuous soil moisture. The VWC test (Sect. 3.1) provides the direct evaluation.

At the seasonal-site mean scale (*n* = 625), total *R*^2^ increased to 0.031. Year remained the dominant predictor (*β* = 0.224, *p* = 0.0002). Precipitation was marginally significant (*β* = 0.133, *p* = 0.037). Temperature became non-significant (*β* = −0.098, *p* = 0.12).

At the annual-site mean scale (*n* = 169), total *R*^2^ reached 0.108. Precipitation and Year were comparable in magnitude (*β* = 0.249, *p* = 0.002 vs. *β* = 0.211, *p* = 0.007). Temperature vanished entirely (*β* = 0.013, *p* = 0.87) (Fig. 3).

**Figure 3.**
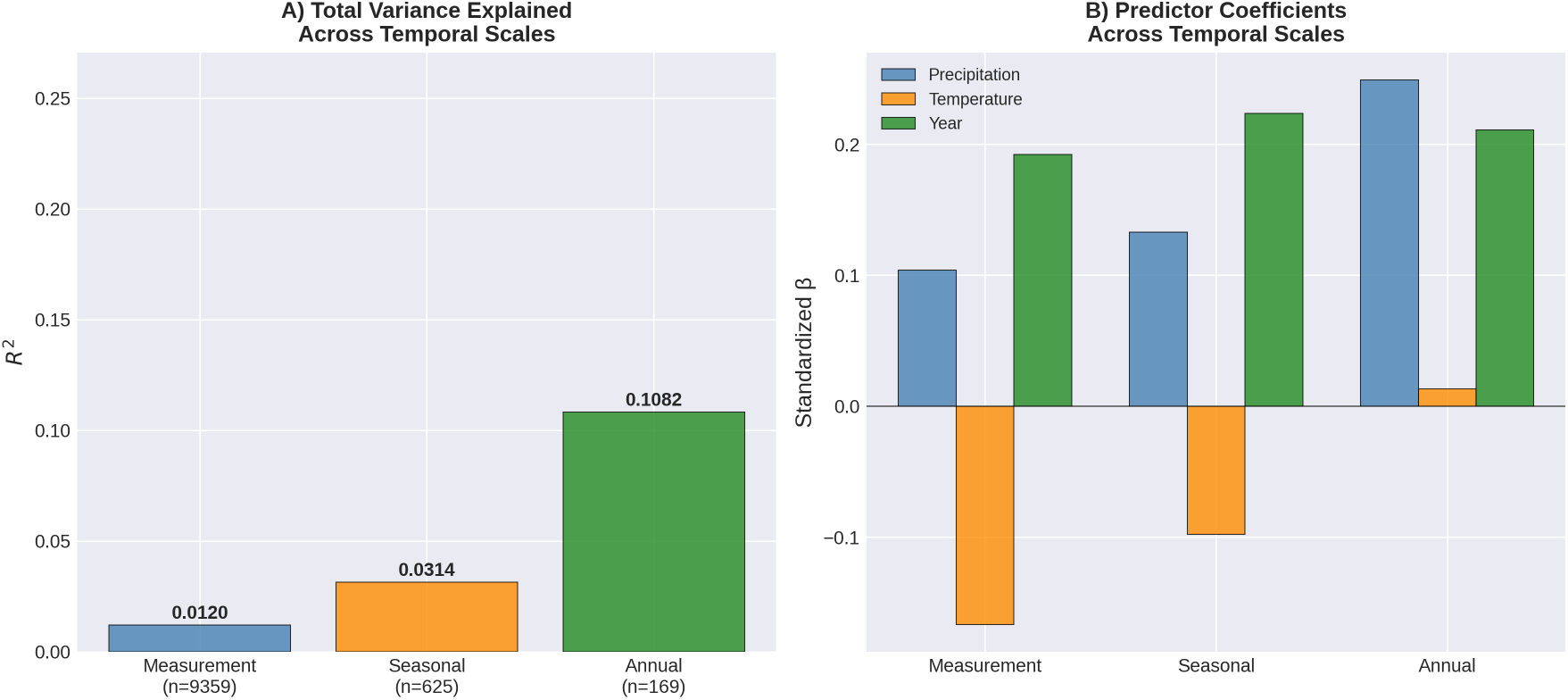
Multi-scale analysis. (A) *R*^2^ across temporal aggregation scales. (B) Standardized predictor coefficients across scales. Even at the annual-site scale, precipitation does not dominate.

Aggregating away measurement noise improved total explanatory power from 1.2% to 10.8%, but it did not produce the pattern predicted by diffusion limitation. At the measurement and seasonal-site scales, Year was the dominant predictor and precipitation contributed the least. At the annual-site scale, precipitation emerged as the strongest individual predictor (*β* = 0.249 vs. *β*_year_ = 0.211): when high-frequency chamber noise is averaged out, interannual precipitation variability does play the leading statistical role in explaining year-to-year flux variation. However, precipitation does not *eliminate* the residual multi-decadal decline, which is absorbed by the Year term (*p* = 0.007). At no scale did Year become non-significant after controlling for moisture, as the diffusion hypothesis predicts. Even under the most generous interpretation (annual site means with all chamber-level variability averaged out), precipitation and a residual temporal trend contribute comparably, and together they explain barely one-tenth of the variance.

### 3.6 Structural breaks at 2002 (BES) and 2011 (HBR)

PELT changepoint detection on annual median CH_4_ flux identified a structural break at 2002 in the BES time series and at 2011 in the HBR time series (Table 5; Fig. 4). These breakpoints were stable across penalty values from 0.05 to 0.5 in the sensitivity analysis.

**Table 5.** Pre- and post-breakpoint flux statistics.

| Site | Breakpoint | Pre-break median<br>(mg C m<sup>-2</sup> h<sup>-1</sup>) | Post-break median<br>(mg C m<sup>-2</sup> h<sup>-1</sup>) | Penalty range<br>(stable) |
| --- | --- | --- | --- | --- |
| BES | 2002 | -1.015 | -0.504 | 0.05–0.5 |
| HBR (WS6-BB) | 2011 | -1.153 | -0.547 | 0.05–0.5 |

*Note:* BES breakpoint precedes HBR by 9 years despite shared regional climate trends.

**Figure 4.**
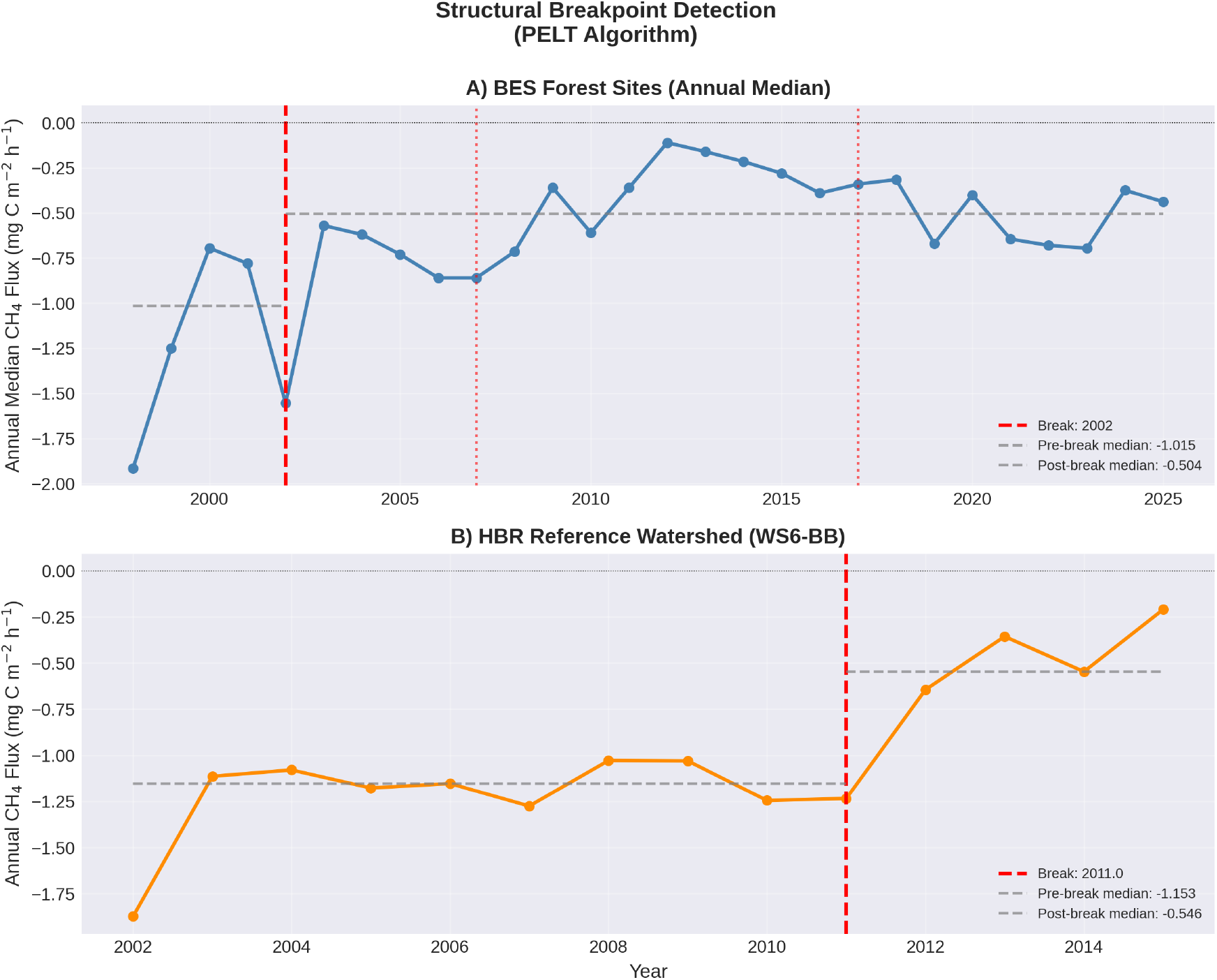
PELT changepoint detection on annual median CH_4_ flux. Structural breaks detected at 2002 (BES) and 2011 (HBR).

The different timing at two independent LTER sites separated by 500 km argues against a common external driver such as regional precipitation change. If increased precipitation drove the decline at both sites, breakpoints should be approximately synchronous or correlated with regional precipitation shifts. The BES break in 2002 precedes the strongest precipitation trends in the mid-Atlantic. The HBR break in 2011 postdates the Ni and Groffman (2018b) dataset by only 5 years and leaves only 4 annual data points in the post-break regime, making it a putative shift rather than a definitive structural break. A sensitivity test dropping the final two years (2014–2015) and re-running PELT on the truncated 2002–2013 record yielded an identical breakpoint across all penalty values tested (0.05–1.0), but the post-break segment in that case consists of just two data points, which is insufficient to establish a distinct statistical regime. We therefore characterize the 2011 HBR break as an emerging shift that requires confirmation through extension of the HBR record. The asynchrony is more consistent with site-specific biogeochemical trajectories than with a shared climatic forcing.

### 3.7 Deposition trends align with flux decline

SO_4_ wet deposition at Hubbard Brook declined from 35.3 to less than 1.0 kg ha^−1^ yr^−1^ between 1978 and 2023, reflecting the cumulative effects of the Clean Air Act and its 1990 Amendments. Inorganic nitrogen 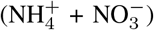 deposition at the Maryland NADP station ranged from 1.2 to 10.5 kg ha^−1^ yr^−1^ over 1999–2023, with a declining trend (Fig. 5).

**Figure 5.**
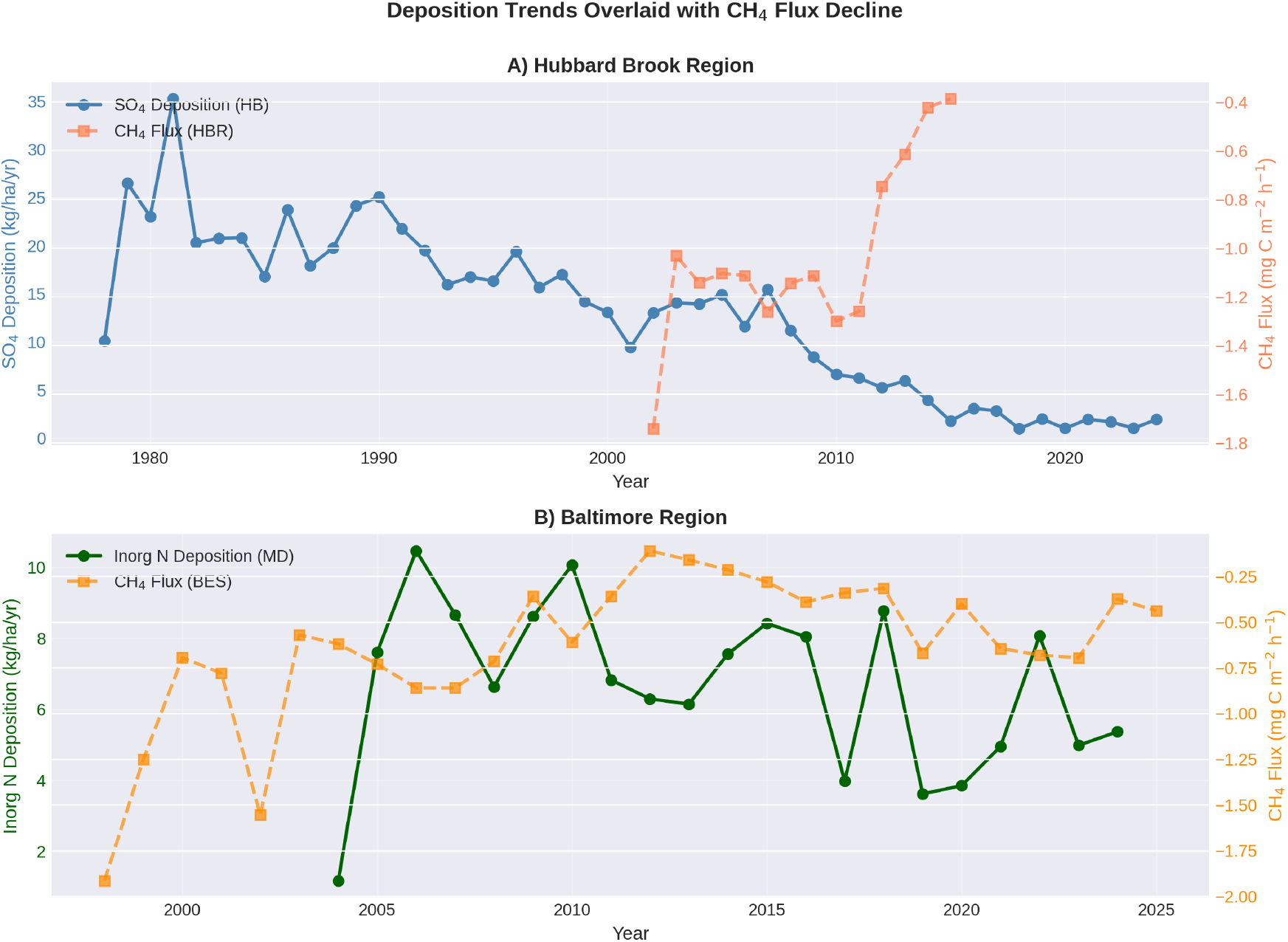
Atmospheric wet deposition trends overlaid with CH_4_ flux time series. SO_4_ deposition at Hubbard Brook (1978–2023) and inorganic N deposition at Beltsville, MD (1999–2023).

The temporal pattern of deposition decline overlaps with the CH_4_ flux breakpoints, though the relationship is indirect. The BES break at 2002 and the HBR break at 2011 both occur while deposition is declining, not increasing. If deposition toxicity were the proximate cause, the break should coincide with peak loading, not with recovery. Two considerations may reconcile this apparent paradox. First, cumulative chronic exposure over 50+ years may cross an irreversibility threshold with a substantial lag, consistent with established nitrogen saturation trajectories in northeastern forests (Aber et al., 1998). Biological collapse due to chronic N-saturation is threshold-dependent and non-linear, which explains why a changepoint algorithm detects a sudden structural break rather than a smooth, gradual decline: the structural break reflects the *manifestation* of accumulated damage to slow-growing methanotrophic populations, not the moment of peak input. Recovery timescales for high-affinity atmospheric methanotrophs are measured in decades to centuries (Lim et al., 2024), so community collapse could become apparent long after peak loading has passed. Second, the decline in acid deposition itself altered soil chemical equilibria (rising pH, shifting base cation ratios, changing aluminum speciation) in ways that may have destabilized microbial communities adapted to decades of acidified conditions. This “recovery disruption” is well documented for other soil processes at Hubbard Brook, where base cation pools continued to decline for decades after atmospheric inputs improved (Likens et al., 1996).

Lysimeter 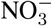 concentrations (1999–2025; *n* = 10,481) showed heterogeneous trends across the three forested BES sites: Hillsdale (urban) exhibited a significant decline (*β* = −0.11 mg L^−1^ yr^−1^, *p <* 10^−26^, *n* = 2,293), consistent with declining atmospheric N inputs; Leakin Park (urban) and Oregon Ridge Upper (rural) showed no significant trend; Oregon Ridge (rural) showed a weak increase (*p* = 0.04, *n* = 1,368). The lack of a coherent network-wide 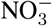 trajectory complicates a simple N-saturation narrative and is consistent with site-specific biogeochemical processing rather than a uniform deposition signal.

This alignment does not establish causation. Deposition, precipitation, temperature, CO_2_, and land use have all changed over the study period. However, the deposition timeline is more consistent with the observed flux patterns than the precipitation timeline, and the urban–rural divergence (Sect. 3.4) argues against atmospheric variables that are spatially uniform across the BES network.

### 3.8 Soil properties

Forest soils at the four characterized BES sites (0–10 cm depth) showed site-level variation in C:N ratio, microbial biomass carbon, net nitrogen mineralization, and net nitrification. This variation documents the biogeochemical heterogeneity underlying the flux patterns and provides context for interpreting site-specific responses. The soil characterization is descriptive; we do not attempt to correlate soil properties with flux trends given the limited spatial replication.

## 4 Discussion

### 4.1 Precipitation-driven diffusion limitation is insufficient at these sites

Five predictions derived from the diffusion limitation framework were tested against 27 years of flux data at BES and 14 years at HBR. Four were not supported: precipitation and direct soil moisture each explained less than 1% of flux variance, no seasonal structure matched diffusion predictions, and urban and rural sites diverged in ways not easily reconciled with shared climate forcing alone (though differences in soil physical properties may contribute). Year dominated the multi-predictor model after controlling for moisture at the measurement and seasonal scales, and remained a significant predictor even at the annual scale where precipitation contributed comparably. The fifth test, the calcium amendment, yielded a null result consistent with the diffusion hypothesis but also with biological alternatives, and constrains the recovery timescale for the methanotrophic community.

Each test alone would be inconclusive. Low *R*^2^ values are typical for individual-measurement regressions in chamber-based trace gas studies (Parkin, 1987), and the calcium null result is consistent with multiple hypotheses. A quadratic moisture–flux model, which accounts for the theoretically unimodal relationship between soil moisture and gas diffusion, did not meaningfully improve explanatory power (Sect. 4.11). However, stratifying at the 2002 changepoint revealed that the pre-breakpoint precipitation–flux relationship was 7× stronger, with a quadratic model explaining 1.8% of pre-breakpoint variance (Sect. 3.1). This pattern is consistent with diffusion playing a minor role in flux variability while the methanotrophic community was intact, and that role becoming undetectable after biological degradation reduced the community’s capacity to respond to moisture fluctuations.

The evidential weight comes from the convergence: five independent tests, using different subsets of the data, different statistical approaches, and different temporal scales, collectively suggest that precipitation-driven diffusion limitation does not adequately explain the multi-decadal decline at these sites. The urban–rural divergence illustrates this: urban sites have stabilized at what we term a “biological floor”—a state in which the high-affinity USC*α* community has been effectively extirpated, leaving only low-capacity methanotrophs that maintain minimal baseline oxidation. If the high-affinity oxidizers are already gone, the sink cannot decline further, regardless of moisture conditions. Rural sites, where USC*α* may persist at reduced abundance, continue to lose capacity. This asymmetry is difficult to account for under the diffusion hypothesis but is consistent with a biological degradation model (Sect. 4.8).

### 4.2 Relationship to Ni and Groffman (2018)

Ni and Groffman (2018b) documented the decline at BES and HBR and proposed increased precipitation as the mechanism. Their analysis established the problem and remains the essential reference for the magnitude and timing of the CH_4_ sink collapse.

This study differs from theirs in five respects. First, the BES record is extended by 9 years (2016–2025), adding substantial information about the post-decline trajectory. Second, direct in-situ VWC data are included, providing the mechanistically relevant variable that Ni and Groffman proxied with precipitation. Third, five independent hypothesis tests are applied rather than a correlation between precipitation trends and flux trends. Fourth, formal changepoint detection with documented penalty calibration is used to identify structural breaks. Fifth, the Hubbard Brook calcium amendment is analyzed as a natural manipulation experiment.

The datasets overlap substantially. The analytical approaches differ. Where Ni and Groffman asked whether precipitation trends correlate with flux trends (they do, weakly), we ask whether diffusion limitation can account for the flux patterns when tested directly. At these sites, over these timescales, it does not.

### 4.3 What does explain the decline?

The five tests do not identify an alternative mechanism. They constrain one. Whatever is driving the decline operates outside the moisture–temperature–diffusion framework, persists after controlling for climatic variables, differs between urban and rural settings, and is not reversed by calcium amendment over 14 years.

The most parsimonious class of explanations is biological: a long-term degradation of the methanotrophic community itself. As noted in the Introduction, the high-affinity USC*α* methanotrophs responsible for atmospheric CH_4_ oxidation are extremely slow-growing, with generation times measured in weeks to months and recovery timescales of decades to centuries (Lim et al., 2024). They are functionally irreplaceable on management-relevant timescales.

### 4.4 Structural diffusion limitation: the earthworm hypothesis

A parallel driver blurs the boundary between “biological” and “physical” mechanisms. Northeastern forests, including parts of the BES and HBR networks, historically lacked native earthworms, and are now subject to invasion by European lumbricids and Asian jumping worms (*Amynthas* spp.) (Bohlen et al., 2004; Chang et al., 2021). Invasive earthworms consume the O-horizon and mix organic matter into the mineral soil, increasing bulk density and destroying macroporosity (Resner et al., 2015). Beyond this physical compaction, the complete removal of the O-horizon eliminates a critical thermal and hydrologic buffer for the mineral soil where methanotrophs reside, exposing the microbial community to amplified temperature and moisture fluctuations that may further degrade oxidation capacity. Because gas-phase diffusivity (*D*_*p*_) is a function of air-filled porosity, earthworm invasion creates a *structural* diffusion limitation (physically reducing *D*_*p*_ through biotic soil compaction) distinct from the *precipitation-driven* (climatic) diffusion limitation posited by Ni and Groffman (2018b). Our analysis addresses the climatic mechanism (increased rainfall filling pore space) and finds it insufficient, but does not address structural changes to the soil matrix itself.

Earthworm invasion also follows an urban-to-rural gradient. Yesilonis et al. (2024) documented shifts in earthworm community composition across the BES urban–rural gradient over two decades, with invasive jumping worms (*Amynthas* spp.) now established at multiple Baltimore sites. This trajectory mirrors the urban–rural divergence in CH_4_ flux (Sect. 3.4): urban sites, where the O-horizon has long been degraded, have stabilized at a biological floor, while rural sites continue to decline as invasion progresses. We do not attempt to partition the relative contributions of nitrogen toxicity and earthworm habitat destruction with the data available here; both mechanisms are consistent with the observed patterns and may act synergistically.

### 4.5 Nitrogen as the primary mechanistic pathway

The literature on nitrogen inhibition of methanotrophy is extensive and mechanistically well characterized. 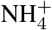 competitively inhibits methane monooxygenase, the enzyme responsible for the initial oxidation of CH_4_ (Schnell and King, 1994; Gulledge and Schimel, 1998). Experimental nitrogen additions reduce forest soil CH_4_ uptake by 33–86% across multiple sites and N forms (Steudler et al., 1989; Bodelier and Laanbroek, 2004; Liu and Greaver, 2009). Long-term fertilization alters the kinetics of atmospheric-concentration oxidation specifically (Gulledge et al., 2004).

Whether ambient deposition levels at BES and HBR are sufficient to cause the observed decline is an open question. Experimental additions in the literature typically exceed ambient deposition by an order of magnitude. Xia et al. (2020) and Cen et al. (2024) demonstrate threshold-dependent responses, with suppression occurring above approximately 25–30 kg N ha^−1^ yr^−1^ in temperate forests. BES inorganic N wet deposition peaked at approximately 10 kg ha^−1^ yr^−1^, below this threshold. However, NADP measures only wet deposition. In urban and peri-urban settings like Baltimore, dry deposition of nitrogen (NO_*x*_ from vehicle emissions, particulate 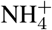) routinely equals or exceeds wet deposition (Lovett et al., 2000). Total N loading at BES was therefore likely 15–25 kg ha^−1^ yr^−1^ historically (consistent with urban-rural deposition gradients documented by Lovett et al., 2000), approaching or exceeding the experimental suppression threshold.

Two additional considerations mitigate the remaining gap. First, BES and HBR have received chronic, continuous deposition for over 50 years; the experimental studies measure responses to acute additions over years to decades. Cumulative chronic exposure may produce effects at lower annual doses than short-term experiments suggest. Second, the relevant comparison may not be current deposition but peak historical loading: SO_4_ deposition at Hubbard Brook reached 35 kg ha^−1^ yr^−1^ in the 1970s, and the legacy effects of that loading on soil chemistry persist decades after atmospheric concentrations declined (Likens et al., 1996, 2002).

### 4.6 The sulfate complication

The role of sulfate in upland methanotrophy is unresolved. Bradford et al. (2001) found that chronic H_2_SO_4_ deposition *increased* CH_4_ oxidation rates in a temperate forest soil, the opposite of what a simple sulfate-toxicity narrative would predict. Su et al. (2025) confirm that the upland sulfate–CH_4_ literature is nascent. The well-developed wetland literature (Gauci et al., 2004; Shen et al., 2025) is not directly transferable to aerated soils where methanotrophy rather than methanogenesis dominates.

We do not claim sulfate deposition as a mechanism for the CH_4_ decline, but the Bradford et al. finding raises a provocative alternative: if chronic acid deposition *stimulated* methanotrophy over the mid-to-late 20th century, then the declines documented at BES and HBR may not represent biological degradation at all, but rather a return to a pre-industrial baseline. Under this “return-to-baseline” hypothesis, the high CH_4_ uptake rates observed in the 1990s were the anomaly—an artifact of decades of acid-stimulated methanotrophic activity—and the post-2002 decline reflects normalization as deposition falls. This interpretation is parsimonious: it requires no irreversible community damage, no nitrogen toxicity threshold, and no earthworm invasion. It predicts that uptake rates should eventually stabilize at a level consistent with pre-industrial soil chemistry, and that sites with the longest deposition histories should show the earliest and largest declines—a pattern broadly consistent with the BES (urban, high cumulative loading, early break) versus HBR (montane, lower cumulative N, later break) comparison.

We cannot distinguish the return-to-baseline hypothesis from the biological degradation hypothesis with the data available here. The key discriminating test would be molecular: if the methanotrophic community is intact but operating at reduced rates because the acid-driven subsidy has ended, USC*α* abundance should remain relatively high. If biological degradation has occurred, USC*α* should be depleted. We note this alternative explicitly because it reframes the narrative: the “decline” may be recovery, viewed from the wrong baseline.

More broadly, deposition trends (including sulfate) show better temporal alignment with flux breakpoints than precipitation does. The mechanistic link, if any, may operate through soil acidification and its cascading effects on base cation availability and microbial community structure (see Brumme and Borken, 1999, who found pH and base saturation explained 77–88 % of among-site variation in CH_4_ oxidation rates across European forests) rather than through direct sulfate toxicity. Resolving this requires experimental work beyond the scope of a flux-based analysis.

### 4.7 Why BES and HBR differ

The two sites broke at different times (2002 vs. 2011), declined at different rates, and operate in different forest types (mixed deciduous urban-rural gradient vs. northern hardwood montane). This heterogeneity is expected under a biogeochemical mechanism and unexpected under a shared climatic driver.

BES soils have received decades of elevated urban nitrogen deposition superimposed on regional acid deposition. HBR soils experienced historically intense sulfate loading (peak *>*35 kg SO_4_ ha^−1^ yr^−1^) followed by rapid decline after the 1990 Amendments. The calcium amendment at WS1 partially reversed acidification effects at HBR but did not restore CH_4_ uptake, consistent with irreversible methanotroph community damage. Borken et al. (2002) documented an analogous non-recovery of CH_4_ uptake at the Solling spruce forest in Germany after more than 10 years of reduced nitrogen and acid inputs.

### 4.8 Recovery is not guaranteed

The post-2012 divergence at BES, with rural forests continuing to decline while urban forests stabilize, suggests that recovery trajectories differ by land use context. Lim et al. (2024) document that high-affinity atmospheric methanotrophs require decades to centuries to recover from persistent disturbance. If USC*α* populations have been depleted by chronic deposition, functional recovery of the forest CH_4_ sink may lag atmospheric improvements by a generation or more, as has been observed for soil base cation pools at Hubbard Brook (Likens et al., 1996).

Urban forests face additional constraints. Edge effects, soil compaction, impervious surface proximity, and heat island effects create conditions that may preclude recolonization by high-affinity methanotrophs even after deposition declines. One interpretation of the urban stabilization is that these sites have reached a biological floor: the high-affinity USC*α* community has been effectively extirpated, leaving only low-affinity, low-capacity methanotrophs that maintain a minimal baseline of CH_4_ oxidation. Under this interpretation, urban flux has stabilized not because conditions have improved, but because there is little remaining biological capacity to lose. Rural sites, where USC*α* populations may persist at reduced abundance, continue to decline as the degradation progresses. This framing generates a testable prediction: urban soils should be depleted in USC*α* relative to rural soils at the same network, detectable through *pmoA* gene surveys (see Sect. 4.11).

### 4.9 Broader context

Brumme and Borken (1999) demonstrated that among-site variation in CH_4_ oxidation across temperate European forests was best explained by soil pH and base saturation (*R*^2^ = 0.77–0.88), not by moisture or temperature. Borken et al. (2002) showed that atmospheric input reductions did not restore CH_4_ uptake at Solling. Täumer et al. (2021) identified soil pH as a key driver of USC*α* abundance and methanotrophic activity. These findings, from independent research groups working on different continents, converge on a common theme: the long-term capacity of forest soils to oxidize atmospheric CH_4_ may be governed primarily by soil biogeochemistry rather than by gas-phase diffusion alone.

The diffusion mechanism likely operates at short timescales (event to seasonal) and controls the high-frequency variability in CH_4_ flux. Wet days produce less uptake than dry days. This is well established. What our data suggest is that diffusion does not control the multi-decadal trend. The patterns are more consistent with a structural shift in the biological capacity of the soil to consume methane than with a dry-to-wet transition.

### 4.10 Limitations

This study has six principal limitations. First, it is based on two LTER networks in the northeastern United States. The conclusions may not generalize to tropical forests, boreal systems, or regions with different deposition histories. Second, we present no direct microbial data. The biological interpretation is inferential, based on the insufficiency of the physical alternative and consistency with the published mechanistic literature. Third, the analysis is observational. Despite the calcium amendment providing a quasi-experimental manipulation, the core flux analyses are correlational. Fourth, precipitation data for urban and rural BES sites are drawn from two PRISM grid cells approximately 20 km apart; any systematic difference between these stations could bias the urban–rural moisture comparison. However, by-site regressions show *R*^2^ *<* 0.07 at every individual site, indicating that station assignment does not mask a real moisture signal within either land-use class. Fifth, chamber-based measurements sample a small fraction of the soil surface and are subject to artifacts (pressure perturbation, collar effects) that eddy covariance methods avoid, though at the cost of sensitivity to the small fluxes involved in atmospheric CH_4_ oxidation. Sixth, the precipitation test uses monthly totals matched to instantaneous chamber measurements; a finer-resolution meteorological proxy such as trailing 7- or 14-day cumulative precipitation might better capture antecedent moisture conditions. The direct VWC test substantially mitigates this concern by measuring the mechanistically relevant variable at the time of flux measurement, but a trailing precipitation index could further refine the climatological test for the pre-2011 period when VWC data are unavailable.

### 4.11 Robustness checks

Thirteen supplemental analyses were conducted to address methodological concerns (Supplementary Information, S1–S13). In brief: (1) removing the *±*3 SD outlier filter *weakened* rather than strengthened the precipitation signal; (2) a quadratic moisture–flux model did not rescue the diffusion signal (*R*^2^ improvement *<*0.05 percentage points); (3) the pre-breakpoint precipitation–flux relationship was 7× stronger than the full-record value but a permutation test could not rule out a selection artifact (*p* = 0.057); (4–5) collar-level random effects (ΔAIC = −24) and AR(1) autocorrelation diagnostics (mean lag-1 *r* = 0.19) confirmed that the Year trend is robust to spatial pseudoreplication and temporal non-independence; (6) a formal interaction test confirmed urban–rural divergence (*p* = 0.007); (7) per-site regressions showed *R*^2^ *<* 0.07 at every individual site, ruling out PRISM station assignment artifacts; (8) the breakpoint permutation test is detailed in the SI; (9) lysimeter NO^−^_3_ trends were heterogeneous across sites (Sect. 3.7); (10–11) the LMM comparison and HBR changepoint sensitivity test (breakpoint stable after dropping 2014–2015) are detailed in the SI; (12) a Precipitation × Post2002 interaction model tested whether the moisture–flux coupling structurally changed at the 2002 breakpoint without subsetting the data; the interaction was not significant (*p* = 0.24), though the Post2002 level shift independently confirmed the PELT breakpoint (*p <* 10^−14^) and Year remained significant after accounting for both (*p* = 0.0004); (13) an LMM with random slopes for Year by site (Year|Site) was fit to address Referee concern that urban and rural sites have fundamentally different temporal trajectories. The fixed Year effect became non-significant (*β* = 0.016, *p* = 0.81) as site-specific slopes absorbed the heterogeneous trends documented in Prediction 4. Critically, the precipitation coefficient was essentially unchanged (*β* = 0.100, *p <* 10^−4^), confirming that the inadequacy of precipitation as a predictor is robust to model specification. The random slopes model confirms that the decline is real but site-specific, consistent with local biogeochemical drivers rather than shared climatic forcing. All thirteen analyses are implemented in `supplemental_robustness.py` in the code repository.

### 4.12 Testable predictions

If the decline reflects degradation of the methanotrophic community, four predictions follow:

1. Archived soils from declining BES and HBR sites should show reduced abundance of USC*α* (high-affinity atmospheric methanotrophs) relative to stable reference sites, detectable through *pmoA* gene surveys and quantitative PCR. Transcriptomic (RNA-based) analysis would be more definitive than DNA surveys alone, since relic DNA from dead or dormant cells can persist in soil for years, inflating apparent abundance.
2. Methanotrophic community composition should differ systematically between urban sites (where the decline has stabilized) and rural sites (where it continues), with urban communities depleted in high-affinity oxidizers.
3. Sites with documented non-recovery of CH_4_ uptake (BES post-2002, HBR post-2011, Solling per Borken et al., 2002) should show convergent community signatures distinct from sites where uptake remains intact.
4. O-horizon thickness and invasive earthworm biomass should correlate with the magnitude of sink decline across BES sites, with urban sites showing thinner O-horizons, higher earthworm density, and lower CH_4_ uptake capacity than rural sites at comparable soil moisture.

These predictions are specific, falsifiable, and addressable with existing molecular ecology methods. Archived soils from BES and HBR, if available, would allow retrospective community analysis spanning the period of decline.

## 5 Conclusions

Five independent tests of the diffusion limitation hypothesis at two long-term ecological research networks, spanning 27 and 14 years of continuous measurement, do not support it as the primary driver of the multi-decadal decline in forest CH_4_ uptake. Neither precipitation nor direct soil moisture explains more than 1% of flux variance. A formally significant urban–rural divergence (*p* = 0.007, interaction test) and a residual temporal trend robust to collar-level pseudoreplication and temporal autocorrelation diagnostics are difficult to reconcile with a purely climatic explanation. The decline documented by Ni and Groffman (2018b) is confirmed and extended with 9 additional years of data, but its attribution to increased precipitation is not supported by the extended record, by direct soil moisture measurements, or by the pattern of asynchronous breakpoints across sites. The timing of those breakpoints, occurring well after peak deposition, is consistent with threshold-dependent nitrogen saturation trajectories (Aber et al., 1998) and the long recovery timescales of high-affinity methanotrophs (Lim et al., 2024). The convergence of these results points toward biological control, consistent with chronic degradation of the methanotrophic community through nitrogen-mediated inhibition, potentially compounded by structural diffusion limitation from invasive earthworm compaction of the soil matrix. Precipitation-driven (climatic) diffusion likely remains an important control on short-timescale flux variability—the pre-breakpoint moisture–flux coupling and its subsequent disappearance support this view—but it does not appear to govern the long-term trajectory of the sink. Confirming the biological mechanism requires molecular evidence from archived soils, and we have outlined four specific predictions to guide that work.

## Code availability

The complete analysis code (`master_analysis.py` and `supplemental_robustness.py`) and data manifest are archived at Zenodo (https://doi.org/10.5281/zenodo.18944402).

## Data availability

All data used in this study are publicly available. BES trace gas fluxes (Groffman and Martel, 2020c), soil moisture and temperature (Groffman and Martel, 2020a), lysimeter chemistry (Groffman and Martel, 2020b), soil properties (Groffman and Raciti, 2020), and permanent plot vegetation surveys (Cary Institute of Ecosystem Studies and Templeton, 2018) are archived at the Environmental Data Initiative (EDI; https://portal.edirepository.org/). HBR CH_4_ flux data are available as knb-lter-hbr.207 (Ni and Groffman, 2018a). Harvard Forest atmospheric CH_4_ concentration data are available as knb-lter-hfr.60 (Crill, 1999). PRISM climate data were obtained from the PRISM Climate Group at Oregon State University (PRISM Climate Group, 2025). NADP wet deposition data were obtained from the National Atmospheric Deposition Program (National Atmospheric Deposition Program, 2025).

## Author contributions

VE designed the study, performed all analyses, and wrote the manuscript.

## Acknowledgements

The Baltimore Ecosystem Study and Hubbard Brook Experimental Forest LTER programs provided the long-term data that made this analysis possible. PRISM climate data were provided by the PRISM Climate Group at Oregon State University. Atmospheric deposition data were provided by the National Atmospheric Deposition Program. All analyses used publicly available data and open-source software. Computational verification of analytical results and preparation of script-generated figures were assisted by Claude (Anthropic). All scientific arguments, data interpretation, and conclusions are the author’s own.

## Appendix A

**Supplementary information: robustness checks**

Thirteen supplemental analyses were conducted to probe methodological concerns and test the robustness of primary results. All analyses are implemented in `supplemental_robustness.py` in the code repository. Results are summarized in `SUPPLEMENTAL_RESULTS.txt`.

### A1 S1: Linear mixed-effects model

The primary multi-predictor model (CH_4_ ∼ Precipitation + Temperature + Year) was fit as a LMM with a random intercept for site, using REML estimation (*n* = 9,359). Site variance (random intercept) was 0.40; residual variance was 5.32. Fixed effects (OLS → LMM): Precipitation *β* = 0.104 → 0.103; Temperature *β* = −0.167 → −0.184; Year *β* = 0.192 → 0.081. The Year coefficient was attenuated in the LMM (part of the OLS temporal trend reflected between-site differences), but remained significant (*p* = 0.003).

### A2 S2: Outlier sensitivity

The precipitation–flux regression and multi-predictor model were re-run without the *±*3 SD per-site-year filter, retaining 126 extreme measurements. Rather than strengthening the diffusion signal, including extreme values *weakened* it: precipitation *R*^2^ dropped from 0.0008 to 0.0003 (*p* = 0.085, non-significant). Multi-predictor *R*^2^ dropped from 0.012 to 0.004. Year remained the dominant predictor (*β* = 0.215, *p <* 10^−5^).

### A3 S3: Non-linear moisture–flux model

A quadratic model (Flux ∼ Precipitation + Precipitation^2^) improved *R*^2^ from 0.077% to 0.12%—a 0.04 percentage point gain. The quadratic term was borderline significant (*p* = 0.044), but total variance explained remained negligible. For direct VWC measurements (*n* = 301), the quadratic term was non-significant (*p* = 0.79) and *R*^2^ remained below 0.03%.

### A4 S4: Pre-breakpoint precipitation regression

Pre-2002 BES data: *R*^2^ = 0.57% (*n* = 1,250, *p* = 0.008), roughly 7× the full-record value. Quadratic 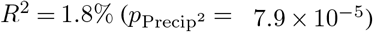. Post-2002: *R*^2^ = 0.01% (*p* = 0.31). Full details in main text Sect. 3.1.

### A5 S5: Urban–rural interaction test

An interaction model (CH_4_ ∼ Year × LandUse) on post-2012 BES chamber data (*n* = 4,639; 2012–2025) yielded a significant Year×LandUse interaction (*β* = −0.071, *p* = 0.007). Secondary analysis on HBR monthly data (2012–2016; *n* = 234): *β* = −0.122, *p* = 0.015. Both confirm statistically divergent trajectories.

### A6 S6: Nested random effects

A collar-level LMM (64 unique Site×Plot×Chamber groups) improved model fit (ΔAIC = −24) and slightly *increased* the Year coefficient (*β* = 0.084, *p* = 0.002) relative to the site-level model (*β* = 0.081, *p* = 0.003). Collar-level pseudoreplication does not attenuate the temporal trend.

### A7 S7: Temporal autocorrelation

LMM residuals showed weak per-site lag-1 autocorrelation (mean *r* = 0.19; Durbin-Watson range 1.34–2.04). This may modestly inflate the Year *p*-value but is unlikely to alter significance at *α* = 0.05.

### A8 S8: Per-site precipitation *R*^**2**^

Individual site regressions: no site exceeded *R*^2^ = 0.07 (max: Cub Hill, *R*^2^ = 0.068). Urban sites mean *R*^2^ = 0.004; rural sites mean *R*^2^ = 0.020. PRISM station assignment does not mask a moisture signal within either land-use class.

### A9 S9: Breakpoint permutation test

A permutation test (*n* = 1,000, shuffling year labels) yielded *p* = 0.057 for the observed Δ*R*^2^ = 0.0056 at the 2002 breakpoint (95th percentile of null: 0.0061). The pre/post coupling difference is suggestive but does not reach formal significance.

### A10 S10: Lysimeter NO^−^_3_ trends

Lysimeter NO^−^_3_ concentrations (1999–2025; *n* = 10,481) at four forested BES sites: Hillsdale (urban) declined significantly (*β* = −0.111 mg L^−1^ yr^−1^, *p <* 10^−26^); Leakin Park (urban) and Oregon Ridge Upper (rural) showed no significant trend; Oregon Ridge (rural) showed a weak increase (*p* = 0.04). Pooled trend was weakly negative (*p <* 10^−4^).

### A11 S11: HBR changepoint sensitivity

Dropping 2014–2015 from the HBR record and re-running PELT on 2002–2013 produced identical breakpoints at every penalty tested (0.05–1.0). At penalties 0.05–0.2, a secondary breakpoint at 2007 appeared in both full and truncated data. The 2011 structural break is not an artifact of the two most extreme post-break observations, though with only 2–4 post-break data points, this putative shift requires confirmation through extension of the HBR record (see Sect. 3.6).

### A12 S12: Precipitation × Post-2002 interaction model

To address the circularity of subsetting the data at the PELT-identified 2002 breakpoint (S4, S9), an interaction model was fit on the full 1998–2025 BES record: CH_4_ ∼ Precipitation + Post2002 + Precipitation × Post2002 (*n* = 9,359). The Post2002 level shift was highly significant (*β* = +0.592, *p <* 10^−14^), independently confirming the PELT breakpoint. However, the interaction term was not significant (*β* = −0.096, *p* = 0.24), indicating that the change in moisture–flux coupling strength at 2002 does not reach statistical significance when tested without data subsetting. Adding Year as a covariate did not alter this result (interaction *p* = 0.27); Year remained significant (*β* = +0.110, *p* = 0.0004), confirming that the secular decline persists after accounting for both the level shift and precipitation. The non-significant interaction is consistent with the permutation test in S9 (*p* = 0.057): the pre/post coupling difference is directionally consistent with a weakening of moisture–flux coupling but does not clear conventional significance thresholds.

### A13 S13: Random slopes LMM

The primary LMM uses a random intercept for site but a fixed (global) slope for Year. Because Prediction 4 demonstrates that urban and rural sites have significantly different post-2012 trajectories, a random-slopes specification (Year|Site) was fit to test whether the fixed Year effect is an artifact of forcing heterogeneous sites to share a common temporal slope. In the random-slopes model, the fixed Year effect became non-significant (*β* = 0.016, *p* = 0.81), as site-specific slopes absorbed the divergent trends. The Year slope variance across sites was 0.026, and the intercept–slope covariance was negative (−0.090), indicating that sites with higher baseline uptake tended to show steeper declines. Critically, the precipitation coefficient was essentially unchanged (*β* = 0.100, *p <* 10^−4^), confirming that the inadequacy of precipitation as a predictor is robust to whether sites are forced to share a common temporal trajectory. The random-slopes model confirms that the decline is real but proceeds at site-specific rates, consistent with local biogeochemical drivers rather than shared climatic forcing.

